# CoGAPS 3: Bayesian non-negative matrix factorization for single-cell analysis with asynchronous updates and sparse data structures

**DOI:** 10.1101/699041

**Authors:** Thomas D. Sherman, Tiger Gao, Elana J. Fertig

## Abstract

**Motivation:** Bayesian factorization methods, including Coordinated Gene Activity in Pattern Sets (CoGAPS), are emerging as powerful analysis tools for single cell data. However, these methods have greater computational costs than their gradient-based counterparts. These costs are often prohibitive for analysis of large single-cell datasets. Many such methods can be run in parallel which enables this limitation to be overcome by running on more powerful hardware. However, the constraints imposed by the prior distributions in CoGAPS limit the applicability of parallelization methods to enhance computational efficiency for single-cell analysis.

**Results:** We upgraded CoGAPS in Version 3 to overcome the computational limitations of Bayesian matrix factorization for single cell data analysis. This software includes a new parallelization framework that is designed around the sequential updating steps of the algorithm to enhance computational efficiency. These algorithmic advances were coupled with new software architecture and sparse data structures to reduce the memory overhead for single-cell data. Altogether, these updates to CoGAPS enhance the efficiency of the algorithm so that it can analyze 1000 times more cells, enabling factorization of large single-cell data sets.

**Availability:** CoGAPS is available as a Bioconductor package and the source code is provided at github.com/FertigLab/CoGAPS. All efficiency updates to enable single-cell analysis available as of version 3.2.

## Introduction

Non-negative matrix factorization (NMF) techniques have emerged as powerful tools to identify the cellular and molecular features that are associated with distinct biological processes from single cell data (Cleary *et al.*, 2017; Zhu *et al.*, 2017; Clark *et al.*, 2019; Welch *et al.*, 2019; Kotliar *et al.*, 2019; Duren *et al.*, 2018). Bayesian factorization approaches can mitigate local optima and leverage prior distributions to encode biological structure in the features (Ochs and Fertig, 2012; Stein-O’Brien *et al.*, 2018). However, the computational cost of implementing these approaches may be prohibitive for large single cell datasets. Many NMF methods can be run in parallel, and thereby leverage the increasing availability of suitable hardware to scale for analysis of large single cell datasets (Schmidt *et al.*, 2009; Ahn *et al.*, 2015; Li *et al.*).

Previously, we developed CoGAPS as a sparse, Bayesian NMF approach for bulk (Ochs and Fertig, 2012; Fertig *et al.*, 2010; Stein-O’Brien *et al.*, 2017) and single-cell genomics analysis (Clark *et al.*, 2019; Stein-O’Brien *et al.*, 2019). CoGAPS was designed to perform Gibbs sampling for a unique prior distribution that adapts the level of sparsity to the distribution of expression values in each gene and cell. While this design allows CoGAPS to adapt to different types of data, it also imposes a constraint on the algorithm that requires the update steps to be proposed sequentially. While the sequential updates of CoGAPS limit implementation of embarrassingly parallel computational approaches, we present a new method for isolating the sequential portion of CoGAPS so that the majority of the algorithm can be run in parallel. Additionally, we derive an optimization for sparse data in order to take advantage of the nature of many single-cell data sets. In combination, these new features in CoGAPS version 3.2 allows for efficient Bayesian NMF analysis of large single cell data sets.

## Approach

### The CoGAPS Algorithm

The input for CoGAPS is a data matrix of single-cell data with N measures (e.g., genes, genomic coordinates, proteins) and M cells, *D* ∈ ℝ^*N*×*M*^, and a number of patterns (features) to learn, *K*. It factors *D* into two lower dimensional matrices, *A* ∈ ℝ^*N*×*K*^) and *P* ∈ ℝ^*K*×*M*^). The columns of the *A* matrix contains relative weights of each measurement for the learned features and the corresponding rows of the *P* matrix contains the relative expression of those features in each cell (Stein-O’Brien *et al.*, 2018). CoGAPS assumes the elements of *D* are normally distributed with mean *AP* and variance proportional to *D*. The algorithm has a Gamma prior on each element of *A* and *P* whose shape hyperparameter has a Poisson prior. This model encodes a sparsity constraint that adapts to the relative sparsity of each gene or cell in the data (Stein-O’Brien *et al.*, 2019). In version 3.2 of CoGAPS, the input matrix *D* can now be passed as either a data matrix or a SingleCellExperiment object. The output for *A* and *P* is stored in a LinearEmbeddingMatrix to enable compatibility with emerging single-cell workflows Bioconductor.

### Asynchronous Updates

Although the algorithm that determines the order of the matrix updates in CoGAPS must be run sequentially, the large number of measurements in genomics data provide feasibility for running the most computationally intensive portion of the algorithm in parallel. Notably, the proposal for which matrix element to update can be made efficiently, whereas evaluating the new value for that element requires an expensive calculation across a row or column of the data. We take advantage of this fact with an asynchronous updating scheme that yields a Markov chain that is equivalent to the one obtained from the standard sequential algorithm (Fertig *et al.*, 2010). In order to do this, we build up a queue of proposals using the sequential algorithm until a conflicting proposal is generated, at which point we evaluate the entire queue of proposals in parallel.

The asynchronous updating scheme heavily relies on the designation of conflicting, or dependent, proposals. Specifically, if two proposals are independent, they must be able to be evaluated in any order without one impacting the sampling distribution of the other. This allows a queue of independent proposals to be evaluated in parallel and still produce a deterministic result. One example of dependent proposals is given here (Figure 1) and a full accounting of all possible conflicts can be found in Supplemental File 1.

**Figure 1.**
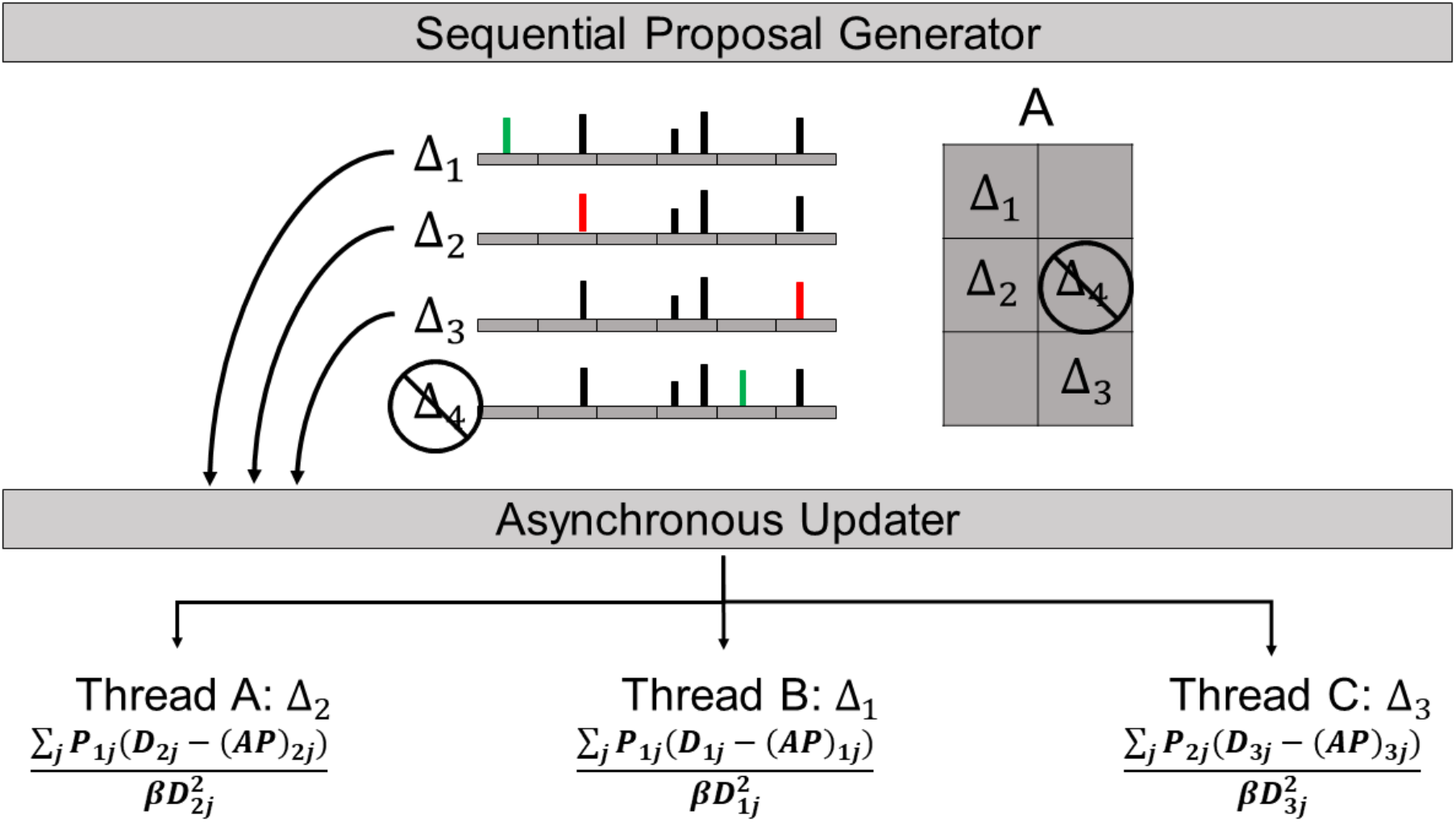
Schematic for the asynchronous updating scheme used in CoGAPS. Updates are proposed sequentially until a conflicting proposal is generated, at which time the existing queue of proposals is evaluated in parallel. In this case, proposed change #4 is conflicting since it is in the same row as #2. When evaluating a proposal, an entire row or column of *AP* is used in the calculation of the conditional distribution used to perform Gibbs sampling. When a change is made in row *n* of *A*, the entire *nth* row of *AP* changes. So, if another change is later proposed in row *n*, the value of *AP* used will depend on the previous proposal thereby changing the conditional distribution for this new proposal. This is exactly the case here for #2 and #4. Changes #1, #2, #3 can be processed in parallel since they do not conflict with each other.

### Sparse Data Structures

Single-cell data tends to be sparse. Therefore, the natural solution for reducing memory overhead is to use sparse data structures to represent the data *D* in the analysis. While the data, *D*, may be very sparse, the weights in *A* and *P* correct for dropout and therefore have a product that may be largely non-zero. Traditionally, CoGAPS caches this product to reduce the number of calculations at each step. However, caching *AP* introduces an unacceptable memory overhead when the data is stored in a sparse format. To address this, we separate the matrix calculations into terms that can be efficiently calculated using only the non-zero entries of *D* and terms that can be precomputed before each batch of updates as described in detail in Supplemental File 1. By doing this, we can make the computation time proportional to the sparsity of the data. However, since storing the data in a sparse format requires the calculation of *AP* during the update steps, there is a performance trade-off that needs to be considered. Typically, when the data is more than 80% sparse it is more efficient to use the sparse optimization, even though it requires calculating *AP*.

### Performance and Benchmarks

We simulated three sparse single-cell datasets with the R package Splatter (Zappia *et al.*, 2017). We varied the level of sparsity in each data set and tested the amount of memory used with and without the sparse optimization enabled. We also measured the run time in both the single-threaded and multi-threaded case. Table 1 gives a high-level overview of the performance differences. For example, with 2,000 genes and 2,000 cells when the data is 90% zeroes, using the sparse optimization and 4 threads will give identical results to the standard algorithm in 1/5 of the time while using 1/25 of the memory.

**Table 1.** Relative performance of the sparse optimization on 2,000 genes and 2,000 cells, baseline is the standard algorithm with 1 thread and no sparse optimization.

| Data Sparsity | Memory (MB) | Runtime (1 thread) | Runtime (4 threads) |
| --- | --- | --- | --- |
| 70% | 0.14 | 1.92 | 0.62 |
| 80% | 0.09 | 1.24 | 0.42 |
| 90% | 0.04 | 0.42 | 0.20 |

We also tested the performance on a single-cell 10X data set generated at the Broad Institute (Bo Li *et al.*). We ran a benchmark on a subset of 37,000 human immune cells (umbilical cord blood) and only kept the 3,000 highest variance genes. In this case, enabling the sparse optimization reduced memory overhead by 82% and cut the run time by 74%. When using 4 threads instead of 1, the run time was cut by an additional 36%.

## Discussion

In this paper, we present an algorithm and software to enable parallelization of CoGAPS to enable analysis of large single cell datasets. This parallelization was done by combining existing methods for Gibbs sampling (Schmidt *et al.*, 2009; Ahn *et al.*, 2015; Li *et al.*) with a new infrastructure for the updating steps in CoGAPS. Prior to the implementation of an asynchronous updating scheme, CoGAPS was applied to large data sets by using a distributed version of the algorithm, GWCoGAPS, that performed analysis across random sets of genes (Stein-O’Brien *et al.*, 2017) or random sets of cells (Stein-O’Brien *et al.*, 2019). This distributed version leveraged the observation that the learned values of *A* and *P* are robust across these random sets. Future work combining the asynchronous and distributed parallelization methods will be critical to further enhance performance by utilizing all CPU cores efficiently.

## Acknowledgments

The authors thank Genevieve Stein-O’Brien, Emily Davis-Marcisak, Loyal Goff, and Ted Liefeld for helpful discussions and benchmarking algorithm performance.

## Supplemental File 1

### 1 The Sequential CoGAPS Algorithm

A description of the sampling steps employed by the serial version of CoGAPS is essential to describe the modifications necessary to enable parallelization. Briefly, CoGAPS is a sparse Bayesian NMF. The input for CoGAPS is a data matrix, *D* ∈ ℝ^*N*×*M*^, an uncertainty matrix, 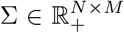, and a number of patterns (latent spaces) to learn, *K*. It factors *D* into two lower dimensional matrices, 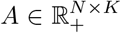 and 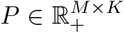. The prior distribution of *A* is given by

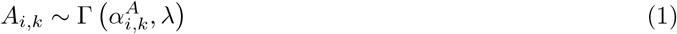

and the prior distribution of *P* is

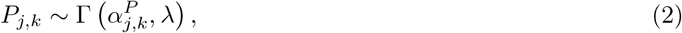

where both 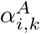 and 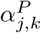 have a Poisson prior with parameter *α*. The likelihood of the data in CoGAPS is given by

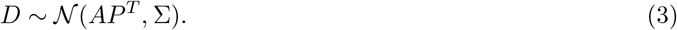

In this model, the values of *α* and *λ* control the relative sparsity of the *A* and *P* matrices. The parameter, *λ*, is set as a function of *α* so that the level of sparsity is inversely proportional to the expected size of each matrix element to better fit the data:

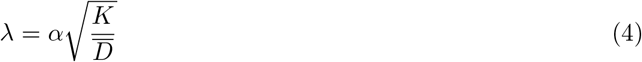

**Figure 1:**
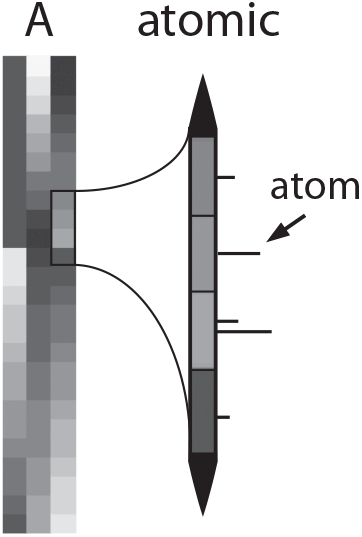
Illustration of the atomic domain for a sample matrix *A*. The values of *A* are depicted in a heatmap in a grayscale where black corresponds to a value of zero (left). The elements *A*_8,3_ to *A*_11,3_ are zoomed in with a depiction of the corresponding bins in 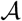, the atomic domain for *A*. Each atom has a position along this domain *l* indicated by the height of the horizontal line segment and magnitude *x* indicated by the length of that line segment. The atomic domain is divided into bins so that the values *l* determine to which matrix element each atom maps. The value of each matrix element is determined by the sum of the magnitudes *x* for all atoms mapping to that element. In this example, *A*_8,3_, *A*_9,3_, and *A*_11,3_ all contain a single atom and whereas *A*_10,3_ contains two atoms. Elements containing no atoms would be exactly zero.

Rather than working with values directly stored in the matrix elements, CoGAPS samples from an atomic prior [1]. This prior is represented by a continuous domain of point masses. Each point mass is called an atom and has a position and mass within this continuous domain (Fig 1). The atomic domain is separated into *N* · *K* bins for the *A* matrix and *M* · *K* bins for the *P* matrix. In this way, each position in the atomic domain maps to a single element of the corresponding matrix. The value of a particular matrix element is equal to the sum of all atom masses in the corresponding bin of the atomic domain or is exactly zero if there are no atoms in the region of the domain corresponding to that matrix element. We let the mass of each atom be distributed according to an exponential prior with parameter *λ*. Because a Gamma distribution is the sum of exponentials, this atomic prior naturally models the Gamma prior imposed on each matrix element. Performing update steps that work directly on the position and mass of the atoms instead of the matrix elements provides a natural way to model the Poisson prior on the shape of the Gamma distribution for each matrix element. It also allows for exact zeros that could not be readily performed sampling from the matrix elements themselves. Moreover, the exponential distribution used for the masses of each matrix element enables Gibbs sampling as described below and derived in detail previously [2, 3, 4, 5].

#### 1.1 Algorithm Overview

Let 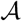 and 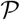 be the atomic domains for the *A* and *P* matrices. Each of the atomic domains is a set of 2-tuples (*l, x*) where *l* ∈ ℕ is the position of the atom and *x* ∈ ℝ_+_ is the mass of the atom (Fig 1). In practice, we restrict the length of the atomic domain so that 0 ≤ *l* ≤ 2^64^. Let 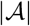 and 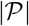 be the number of atoms in each domain. We also define 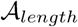 and 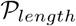 to be the size, or number of available positions, of each domain. The size is equal to the closest number to 2^64^ that the number of bins, or matrix elements, evenly divides. This allows each atom to be uniformly distributed and each matrix element to have an equal sized portion of the atomic domain mapping to it.

As shown in Algorithm 1, update steps are run in batches, alternating between the *A* and *P* matrix. An iteration of CoGAPS is defined as a batch of updates for each the *A* and *P* matrix. The algorithm runs in two phases, equilibration and sampling. Each of these phases is run for a pre-specified number of iterations, input by the user. The goal is to pick a number of iterations large enough for the sampling phase to obtain samples from the posterior distribution, *p*(*A, P*|*D, σ*). These samples are averaged to get the final values for *A* and *P* as the Bayes minimum mean square error estimator.

**Algorithm 1:**
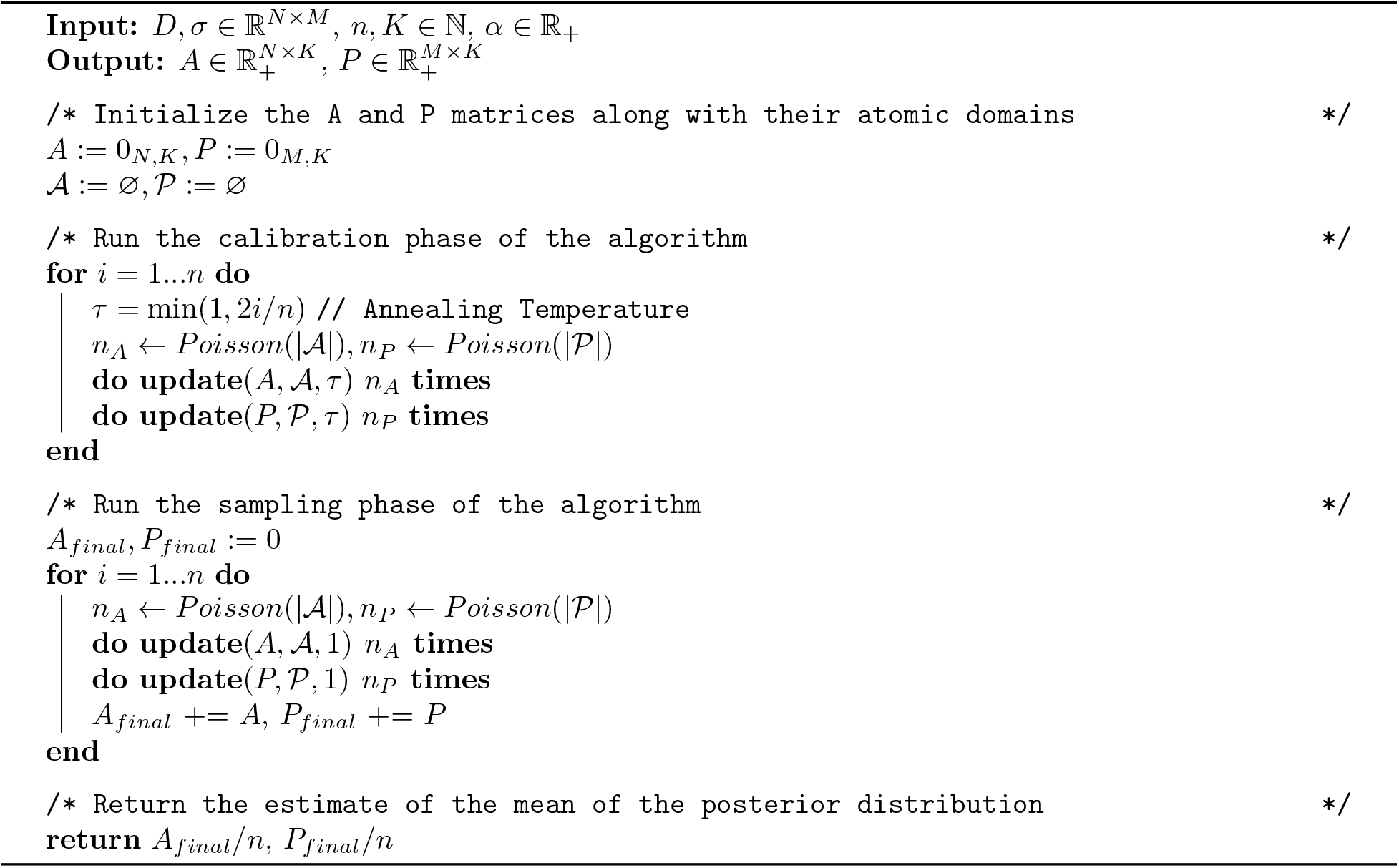
CoGAPS

When updating the *A* and *P* matrices, CoGAPS preforms sampling steps to directly update the values of the atoms in 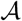 and 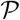. There are four types of updates that can happen in the atomic domain: 1) Birth - creates a new atom 2) Death - destroys an existing atom 3) Move - changes the location of an existing atom 4) Exchange - takes a portion of the mass from one atom and combines it with another. These four steps are selected as the updates for the atomic domain because they preserve the desired prior distributions of the matrix elements and enable Gibbs sampling. Each update step starts by randomly selecting one of these 4 changes. In any given update step, either one or two matrix elements will change. In the Bayesian NMF algorithm presented in [6], Gibbs sampling is done by sampling from the conditional posterior distribution of the matrix element, i.e. from *p*(*A*_*i,j*_|*A*_\(*i,j*)_, *P, D, σ*) or *p*(*P*_*i,j*_|*P*_\(*i,j*)_*, A, D, σ*). Due to the atomic domain in CoGAPS, sampling is instead done on the atoms themselves, i.e. sampling is done from *p*(Δ*A*_*i,j*_|*A, P, D, σ*) or *p*(Δ*P*_*i,j*_|*P, A, D, σ*). As described above, the sparsity of the matrix elements is determined by the Poission prior with parameter *α*. This distribution can be sampled from by altering the relative probability of a Birth or a Death step according to the sampling steps in Algorithm 2.

**Algorithm 2:**
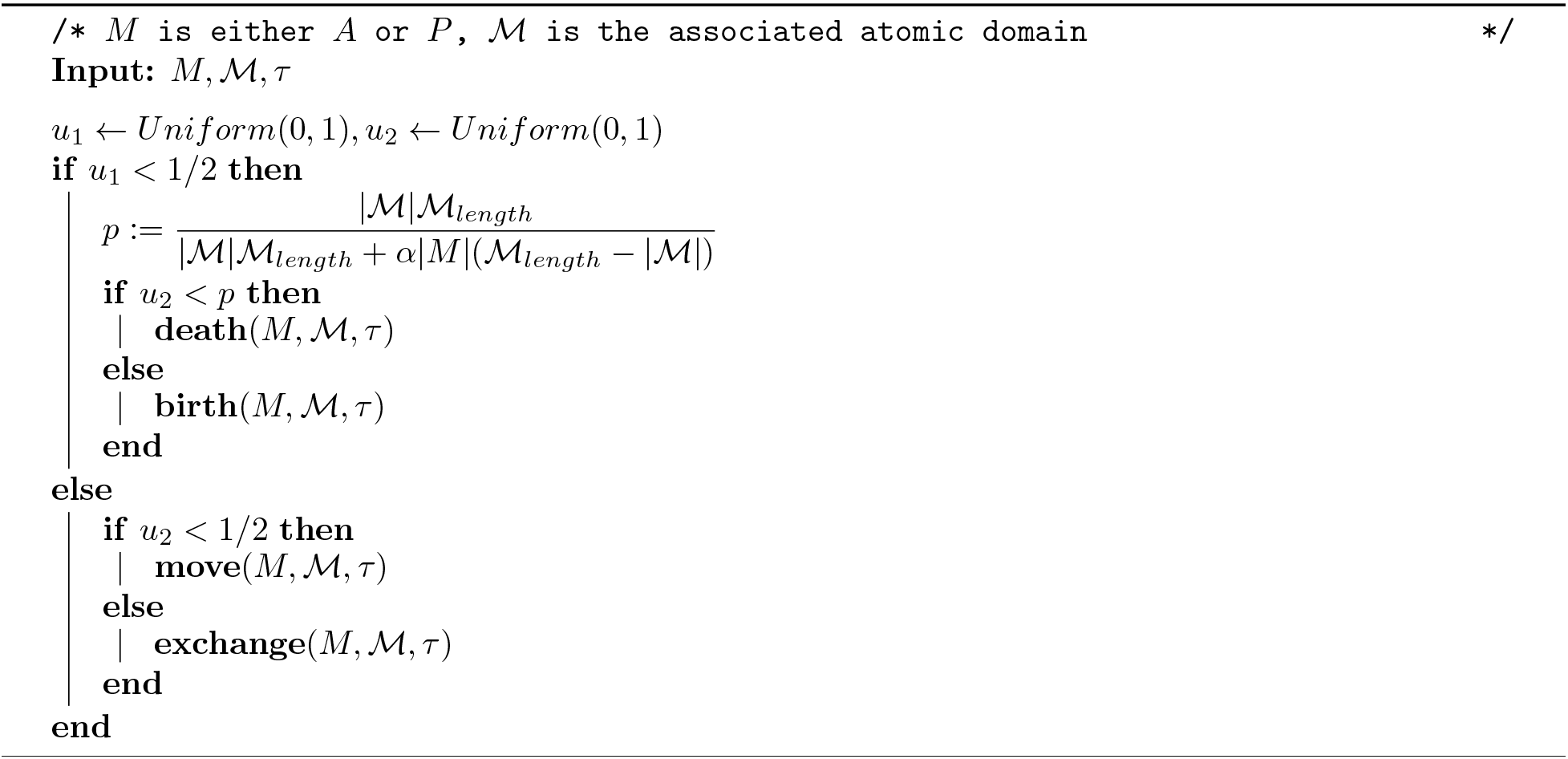
Update

#### 1.2 The Update Steps

As we describe above, there are four types of updates to the atomic domain in CoGAPS: birth, death, move, and exchange. The probability of an update step being any of these is described in Algorithms 1 and 2. In this section, we will describe how each step works. We break down each step into two parts: the proposal and the evaluation. This distinction becomes important for when we present the asynchronous updating scheme in the next section. The proposal phase determines which atoms are affected and the evaluation phase determines how large the effect is, and whether or not to accept certain changes. Note that we present the following math in terms of *M* and 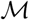 for the matrix and atomic domain, since these steps are the same for either the *A* or the *P* matrix. We also reference quantities *s* and *s*_*μ*_ which are defined in the Appendix, with full derivations available in previous work [5].

##### 1.2.1 Birth

In this type of update we are creating a new atom in a random location of the atomic domain.

###### Proposal

The proposal part of this step involves selecting a random, open location for the atom.

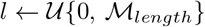 such that 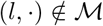

Let *l* ⟼ *r, q*, where *r, q* are the corresponding row and column in the matrix.

###### Evaluation

We want to sample a mass for this atom *x*. This corresponds to a change in the *M_r,q_* element, i.e. Δ*M*_*r,q*_ = *x* which is exponentially distributed. We want to sample *x* from *p*(Δ*M*_*r,q*_|*A, P, D, σ*). Previously, we have shown that under the normal likelihood this distribution follows a truncated normal with the following parameters:

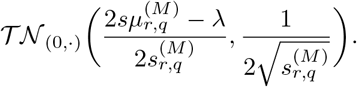

We sample *x* from this distribution and add the new atom, (*l, x*), to 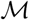.

##### 1.2.2 Death

In this type of update, we select an atom at random and remove it from the atomic domain. In practice, we take an additional step that helps the algorithm converge faster. Rather than just removing the atom outright, we give it a chance to re-sample a new mass and then keep that atom with some probability based on how the new mass effects the likelihood, *p*(*D*|*A, P, σ*).

###### Proposal

In order to propose a death step, we only need to select an atom uniformly at random. Let (*l, x*_0_) be the location and mass of this atom, and let *l* ⟼ *r, q* where *r, q* are the corresponding row and column of the matrix.

###### Evaluation

When evaluating the death step, we need to first remove the atom from the atomic domain and then sample a new mass in the same location. Rather than actually removing the mass and updating the matrix, we can instead directly compute *p*(Δ*M*_*r,q*_|*x*_0_*, A, P, D, σ*) where *x*_0_ is the original mass that has been removed. This distribution is a truncated normal with the following parameters:

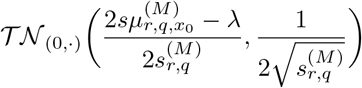

We sample *x* from this distribution to get a new atom, (*l, x*). However, we only add it to 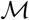 with some probability. Otherwise, the atom is removed from the atomic domain completely. The probability of keeping this new atom is 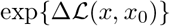, where 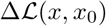 is defined as the log-likelihood ratio:

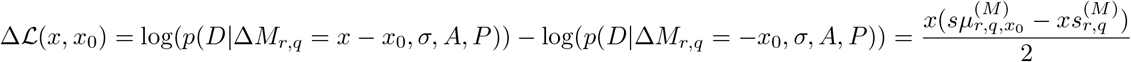

##### 1.2.3 Move

In this type of update, we are moving an atom from one location to another. We restrict the move so that the atomic domain is not re-ordered, i.e. an atom can only move between it’s left and right neighbors.

###### Proposal

We select an atom uniformly at random. Let (*l*_0_*, x*) be the location and mass of this atom. Let *l*_*left*_ and *l*_*right*_ be the locations of the left and right neighbors of this atom. If the neighbor does not exist then *l*_*left*_ = 0 and 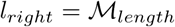. We then sample a new location *l* from the uniform distribution:

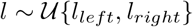

Let *l*_0_ ⟼ *r*_0_*, q*_0_ and *l* ⟼ *r, q* be the corresponding row and column each location maps to.

###### Evaluation

If *r*_0_ = *r* and *q*_0_ = *q*:

Automatically accept the change as there is no effect on the matrix, simply update the atom’s position.

If *r*_0_ ≠ *r* or *q*_0_ ≠ *q*:

Accept the move with probability, exp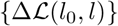, where 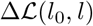 is defined as the log-likelihood ratio:

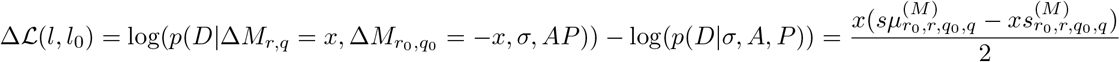

##### 1.2.4 Exchange

In this type of update, we are exchanging mass between two atoms. First an atom is selected at random. Then we use a Gibbs sampling approach to determine how much mass to exchange between this atom and its right neighbor.

###### Proposal

We select an atom uniformly at random. Let (*l*_1_*, x*_1_) be this atom and let (*l*_2_*, x*_2_) be its right neighbor. If the right neighbor does not exist (i.e. *l*_1_ is the right-most atom) then set the right neighbor equal to the left-most atom. Let *l*_1_ ⟼ *r*_1_*, q*_1_ and *l*_2_ ⟼ *r*_2_*, q*_2_ be the corresponding row and column each location maps to.

###### Evaluation

If *r*_1_ ≠ *r*_2_ or *q*_1_ ≠ *q*_2_:

We want to sample the mass increase to the first atom, let this be *x* (i.e. Δ*x*_1_ = *x*). Since we are exchanging mass, we must preserve the total mass between both atoms. To accomplish this we must have *x* ∈ (−*x*_1_*, x*_2_). At the end of this step we will set the new masses to be *x*_1_ ⟶ *x*_1_ + *x* and *x*_2_ ⟶ *x*_2_ − *x*. We want to sample *x* from 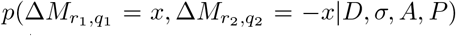. This distribution is a truncated normal with the following parameters:

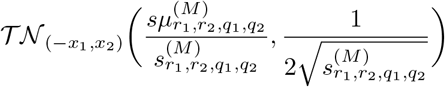

If *r*_1_ = *r*_2_ and *q*_1_ = *q*_2_:

In this case, there is no change to the matrix elements or the structure of the atomic domain so it is safe to ignore this update.

#### 1.3 Annealing Temperature

When the algorithm is in the equilibration phase, we use a strategy known as simulated annealing. This allows the initial samples to be dominated by the prior distributions rather than the likelihood. Let *τ* be the temperature, then in all cases we have *s* ⟵ *τs* and *sμ* ⟵ *τsμ*. Eventually we reach *τ* = 1, in which case *s* and *sμ* are calculated exactly as described above.

### 2 Asynchronous Update Steps

The previous section outlined the standard CoGAPS algorithm and showed how the update steps of the algorithm work in detail. It is important to note that, in the standard algorithm, these update steps must be run sequentially. The proposal phase of each update step requires knowledge about the current state of the atomic domain. If the updates were run in parallel, it would not be possible to know the current state of the domain and the algorithm would be fundamentally different. The evaluation phase, however, is not so restricted. This phase is just a large matrix calculation. If two update steps involve unrelated segments of the matrices, then there is no reason the evaluation phase cannot be run in parallel. Conveniently, the proposal phase is relatively inexpensive to compute when compared to the evaluation phase. This concept is the foundation of the asynchronous version of CoGAPS. The proposal phase of the update steps is run sequentially in order to build up a queue of “independent” proposals which are then evaluated in parallel.

The details of the asynchronous algorithm come down to defining independent proposals. We want two proposals to be independent only if they can be evaluated in any order without changing the state of the algorithm (i.e. the atomic domain and matrix values). If we are able to do that, then the algorithm is straightforward: propose as many updates as possible until there is some dependency, then evaluate all the independent proposals in parallel. If our definition is correct, this will result in the same state of the algorithm as if we processed everything sequentially. Since it is difficult to define what makes proposals independent, we will instead make an exhaustive list of the conflicts that could prevent us from adding a new proposal to the queue. The next three sections explore these conflicts.

#### 2.1 Conflict in Matrix Calculations

The first type of conflict arises in the calculations of *s* and *sμ*. Both of the quantities depend on the values in the *A* and *P* matrices. For the sake of example, say that we are proposing a change to *A*_*r,q*_. In this case, we will calculate 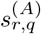 and 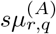 which both depend of the *r*^*th*^ row of *AP*^*T*^. If we accept a change to *A*_*r,q*_ we will also be changing the entire *r*^*th*^ row of *AP*^*T*^ and this is where the conflict happens. If there are two proposals in the same row of *A*, then the outcome of one will effect the calculations of *s* and *sμ* for the other. Therefore, any two proposals that effect the same rows are not independent. This observation has been made in other matrix factorization algorithms and similiar schemes were designed in order to run the factorization parallel [6, 7, 8, 9]. To check for this conflict, we keep a table of all the rows that are currently used by proposals in the queue. If any new proposal has an overlap, we stop the queue and move on to the evaluation phase. Surprisingly, this conflict accounts for nearly all the possible conflicts that could arise, however there are some corner cases described in the next section.

#### 2.2 Atomic Domain Ordering and Finding Neighbors

Checking for a matrix conflict will take care of most of the conflicts that can occur between proposals, however there is one type of update that can be problematic without causing any matrix conflicts. When doing a move step, we select an atom at random and move it somewhere between its neighbors. That means we must be able to calculate the positions of both neighbors. Most of the time, neighboring atoms will map to matrix elements in the same row, but this is not guaranteed to be the case - especially since the atomic domain can be very sparse. Therefore, it is possible that the starting location and proposed new location for a moved atom could map to matrix elements in a different row than the neighboring atoms. In this case, it would be possible for the neighbors to be involved in a previously queued update step but not have any conflicts in the matrix calculations. To account for this, during a move step we must explicitly check that the neighboring atoms are not involved in any previously queued updates.

There is additional type of conflict that can arise directly in the atomic domain during the birth step. Any queued death step can open up more locations for future atoms to be created in. The probability of a location chosen at random being already occupied is astronomically small (in practice the atomic domain has 2^64^ possible locations). The chance that not only was this location occupied, but the atom also was removed in the last handful of updates is so small that we ignore this conflict. New atoms are created among all the available locations, not counting ones that could be made available from the current queue.

#### 2.3 Birth/Death Decision

Direct conflicts between update steps are not the only way that the queue can be stopped. It is possible to run into an issue before we even decide what type of update to queue next. When deciding between a birth and death step, we need to know the current size of the atomic domain. This allows the algorithm to enforce a sparsity constraint by tightly controlling the total number of atoms. Every time a proposed update is queued, it becomes less clear what the total number of atoms will be. At update *t* if we know there *n* atoms, and we propose a death step, then at *t* + 1 there are between *n* − 1 and *n* atoms. The more updates that are queued up, the larger this range becomes.

Fortunately, the function that determines the probability of death vs birth is a monotonic function of the total number of atoms. Let *P*_*death*_(*n*) be the death probability when there are *n* atoms and let *n*_*t*_ be the total number of atoms at time *t*. If we have evaluated all updates up to time *t* then we know *n*_*t*_ exactly. If we propose *k* more updates however, we do not know what *n*_*t*+*k*_ is. Based on the types of the proposed updates we can calculate a range for this value. Let *N*_*deaths*_ and *N*_*births*_ be the number of deaths and births proposed in the current queue. Then we know that *n*_*t*_ − *N*_*deaths*_ + *N*_*births*_ ≤ *n*_*t*+*k*_ ≤ *n*_*t*_ + *N*_*births*_. We therefore know that *P*_*death*_(*n*_*t*_ − *N*_*deaths*_ + *N*_*births*_) ≤ *P*_*death*_(*n*_*t*+*k*_) ≤ *P*_*death*_(*n*_*t*_ + *N*_*births*_).

When making the decision between a birth step and a death step at time *t*, we draw 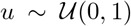. If *u* < *P*_*death*_(*n*_*t*_) we propose a death step, otherwise a birth step. For making this decision *k* proposals into the future, we have three possible outcomes instead of two. If *u* < *P*_*death*_(*n*_*t*_ − *N*_*deaths*_ + *N*_*births*_) then we know we have a death step. If *u* > *P*_*death*_(*n*_*t*_ + *N*_*births*_) then we know we have a birth step. If *u* is between these two values then we cannot make a decision and the result is indeterminate. We must first evaluate the current queue in order to get the exact value for *n*_*t*+*k*_. So, in this case the queue stops being populated and we move on to the evaluation phase.

### 3 Sparse Data Optimization

The previous sections have discussed strategies improving the run time of CoGAPS, but have not addressed the amount of memory used by the algorithm. Memory overhead is one of the most significant problems facing the analysis of large single-cell datasets. Fortunately, these datasets tend to be extremely sparse and so there is an opportunity for reducing the memory overhead of the algorithm by storing the data in sparse data structures. Computing the values of *s* and *sμ* on sparse data structures however is not straightforward. It is necessary to derive a sparse-friendly version of the *s* and *sμ* calculations to avoid increasing run time in solving for memory usage.

#### 3.1 Sparse Matrix Calculations

In this case we will derive the equations just for the *A* matrix, since the derivation for the *P* matrix is identical except for the labels. We first define the non-zero indices of a given row of the data as *γ* and make an assumption about the standard deviation, *σ*:

Let *γ*(*i*) = {*j*: *D*_*ij*_ > 0} and 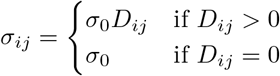

In the Appendix we show the full derivation of the following identities:

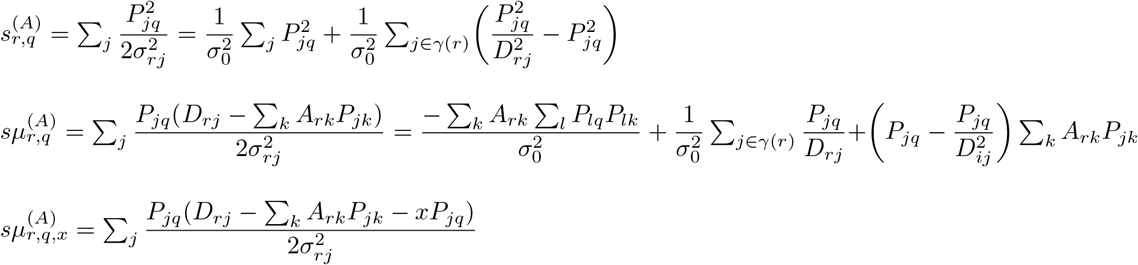

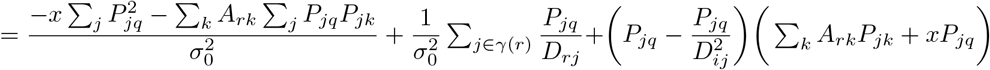

(This is for *r* = *r*_1_ = *r*_2_, the *r*_1_ ≠ *r*_2_ case requires no additional derivation)

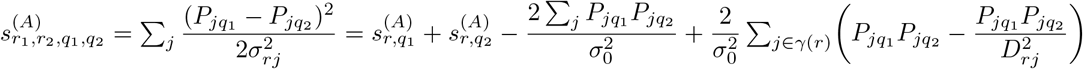

Note that we can longer use the *AP*^*T*^ matrix in the calculations since it is not feasible to cache it when working with large, sparse datasets. Even when *D* is very sparse, the product of *A* and *P* will not be. In fact, *AP*^*T*^ is nearly all non-zero is most real world cases. Storing this large of a matrix defeats the entire purpose of the optimization, so we must compute *AP*^*T*^ on the fly.

#### 3.2 Runtime Complexity

Each of these calculations follow a similar pattern: there is some initial term and then a sum over the non-zero indices of the data. Without the optimization, we would have to sum over all the indices of the data. Therefore, as long as the initial term doesn’t take too long to compute, the full computation should be faster than the standard computation by a factor of the data sparsity. In this section, we will explore the runtime complexity and computational strategies for the initial terms.

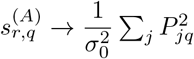

This term only depends on the *P* matrix which is constant while *A* is being updated, so it can be precomputed ahead of a batch update step for *A*. To compute this term for all values of *q* requires *O*(*MK*) time where *M* is the number of samples and *K* is the number of patterns. A batch of updates can be done to *A* is *O*(*NMK*) time where *N* is the number of features. Thus creating a lookup table for this initial term has a negligible cost when compared to the full batch of updates. Once we have a lookup table, then computing the initial term takes *O*(1).

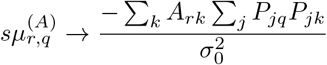

The Σ_*j*_*P*_*jq*_*P*_*jk*_ term only depends on *P* so we can also make a lookup table for this value. Creating this lookup table takes *O*(*MK*^2^) time, whereas the full batch of updates takes *O*(*MNK*). In practice, *K* ≪ *N* so this is an acceptable overhead to incur. Once we have a lookup table, the initial term takes *O*(*K*) to compute, which makes the update step go from *O*(*MK*) to *O*(*MK* +*K*) which again is acceptable since *K* ≪ *M*

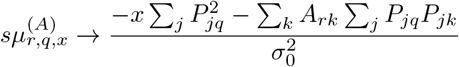

We have already created a lookup table for both 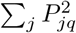 and Σ_*j*_*P*_*jq*_*P*_*jk*_ so there is no additional cost for this step, and the remaining computation takes the same amount of time as the previous quantity, 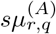, so no additional work needs to be done to show this incurs an acceptable overhead.

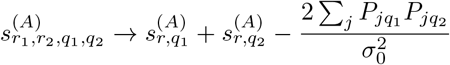

While it may seem that 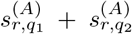 are part of the initial term, both of these quantities can be computed alongside the main term, so no additional run time cost is incurred. The 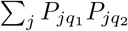 term has already been addressed above, so there is no additional work needed.

### 4 Appendix

#### 4.1 Single Matrix Element Calculations

These calculations are relevant when a single matrix element is being changed (such as in birth or death). Let *r, q* be the row and column of the changed element and let ^(*A*)^ or ^(*P*)^ denote which matrix is being changed.

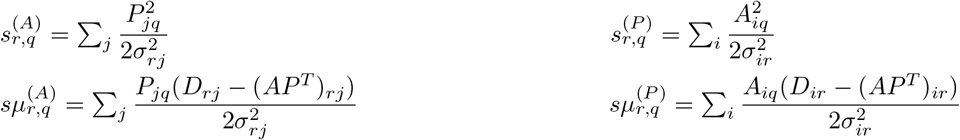

In this case, we want to make the same *sμ* calculation, but with the assumption that a change *x* has already been made to the (*r, q*) matrix element.

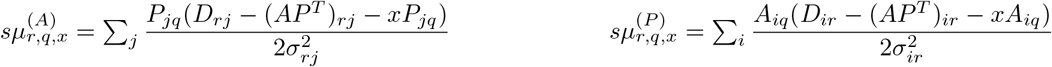

#### 4.2 Multiple Matrix Element Calculations

These calculations are relevant when two matrix elements are being changed at the same time (such as in move or exchange). Let *r*_1_*, q*_1_ be the row and column of the first element and *r*_2_*, q*_2_ be the second element.

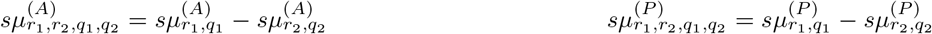

IF *r*_1_ = *r*_2_:

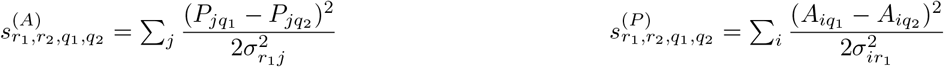

If *r*_1_ ≠ *r*_2_:

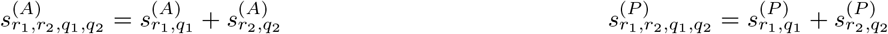

#### 4.3 Sparse Matrix Calculations

Let *γ*(*i*) = {*j*: *D*_*ij*_ > 0} and 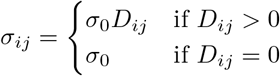

##### 4.3.1 *s*_*r,q*_

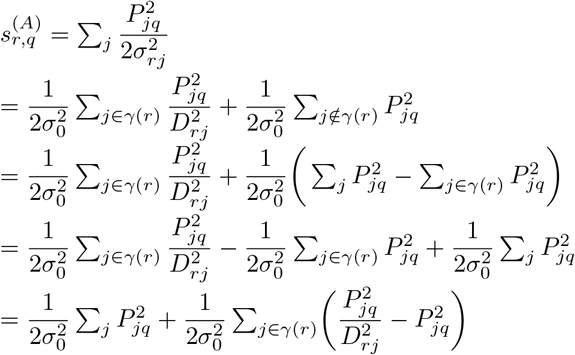

##### 4.3.2 *sμ*_*r,q*_

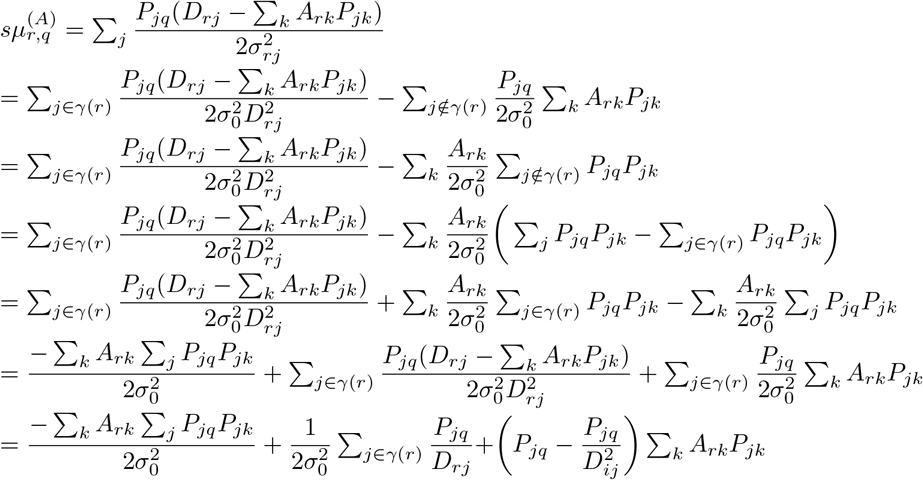

##### 4.3.3 *sμ*_*r,q,x*_

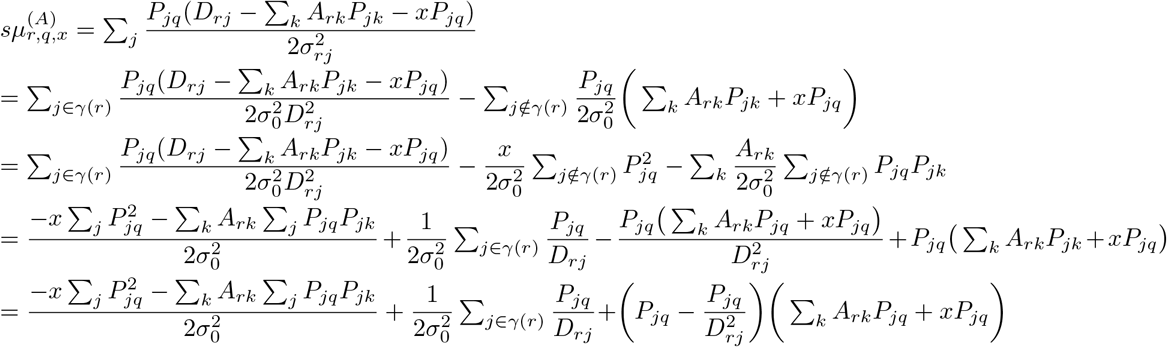

##### 4.3.4 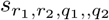

In this case, *r* = *r*_1_ = *r*_2_

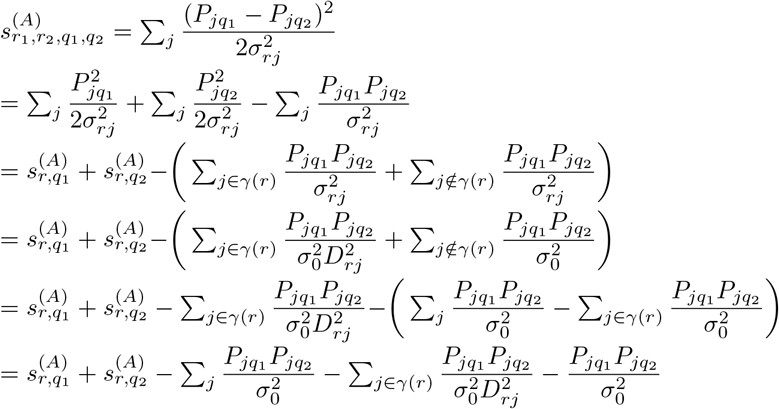

#### 4.4 Sparse *χ*^2^ Calculation

The following derivation shows a more sparse-friendly way of calculating *χ*^2^. This isn’t relevant for the updating portion of the algorithm, but the performance is somewhat relevant since it is calculated periodically throughout the run of the algorithm.

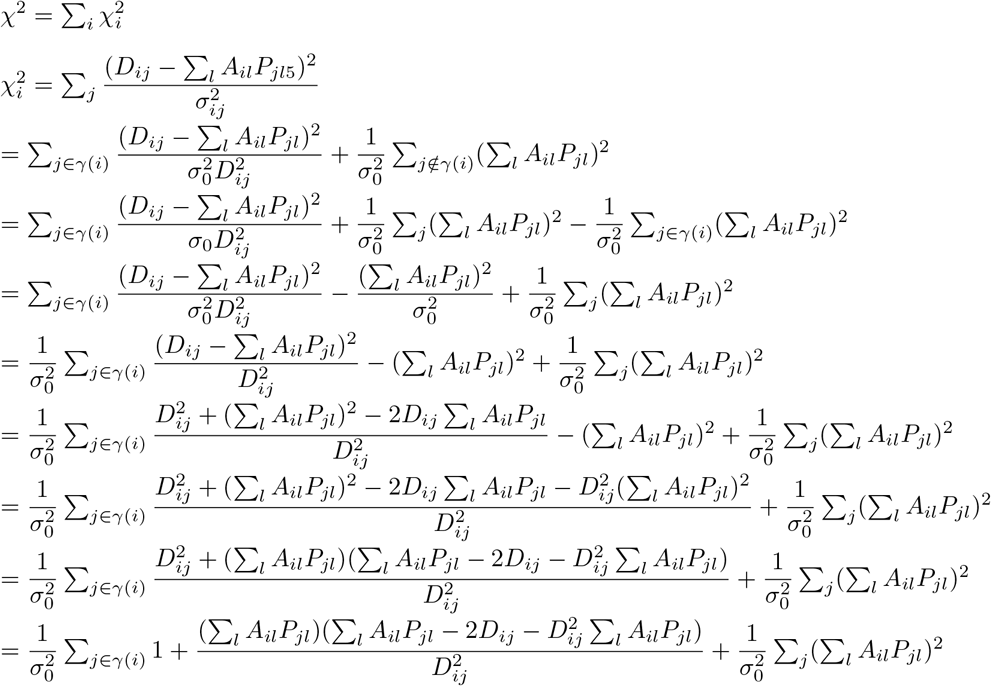

